# SeeCiTe: a method to assess CNV calls from SNP arrays using trio data

**DOI:** 10.1101/2020.09.28.316372

**Authors:** Ksenia Lavrichenko, Øyvind Helgeland, Pål R Njølstad, Inge Jonassen, Stefan Johansson

## Abstract

**Motivation:** Single nucleotide polymorphism (SNP) genotyping arrays remain an attractive platform for assaying copy number variants (CNVs) in large population-wide cohorts. However current tools for calling CNVs are still prone to extensive false positive calls when applied to biobank scale arrays. Moreover, there is a lack of methods exploiting cohorts with trios available (e.g. nuclear family) to assist in quality control and downstream analyses following the calling.

**Results:** We developed SeeCiTe (Seeing Cnvs in Trios), a novel CNV quality control tool that post-processes output from current CNV calling tools exploiting child-parent trio data to classify calls in quality categories and provide a set of visualizations for each putative CNV call in the offspring. We apply it to the Norwegian Mother, Father, and Child Cohort Study (MoBa) and show that SeeCiTe improves the specificity and sensitivity compared to the common empiric filtering strategies. To our knowledge it is the first tool that utilizes probe-level CNV data in trios to systematically highlight potential artefacts and visualize signal intensities in a streamlined fashion suitable for biobank scale studies.

**Availability and Implementation:** The software is implemented in R with the source code freely available at https://github.com/aksenia/SeeCiTe.

## Introduction

The term copy number variant (CNV) commonly refers to a segment of DNA (typically >1kb in size) that varies among individuals in the number of copies. There is an established and ever-growing evidence of significant role that CNVs (and other types of structural genomic variation) play in disease and evolution (Bailey and Eichler (2006), Feuk, Carson, and Scherer (2006), Zarrei, MacDonald, Merico, and Scherer (2015), Girirajan, Campbell, and Eichler (2011)).

Among the technologies employed to assay genomic data for variation in tens of thousands of individuals, SNP genotyping arrays remain attractive due to their permissive cost and established methodology. Importantly, SNP arrays data already exists for a number of population-wide cohorts combined with appropriate tools allow for detection of CNVs. Notable examples are the UK Biobank (Kendall et al. (2017)) and the Norwegian Mother, Father, and Child Cohort Study (Magnus et al. (2016)). These projects were often designed with focus on SNPs and thus were assayed on arrays that may pose difficulties in consequent CNV calling (Pinto et al. (2011)).

For this reason, a plethora of methods have been developed in the past decades that may be utilized for CNV calling in the data produced using various array platforms and chip designs. Among the most often used are PennCNV (Wang et al. (2007)) (1398 citations at the time of the writing), QuantiSNP (Colella et al. (2007)) (580 citations), cnvPartition (Illumina proprietary software) and Birdsuite (Affymetrix proprietary software). Common to all existing methods are persisting issues with precision in CNV calling. Array-based CNV calling methods are known to have false positive rates as high as 24% (PennCNV, Eckel-Passow, Atkinson, Maharjan, Kardia, and de Andrade (2011)), and inter-software reproducibility below 50% (Pinto et al. (2011)). This led in turn to numerous efforts and strategies aimed at increasing the specificity of a given CNV calling pipeline (Zhang et al. (2014), Li et al. (2018)), for example, involving visual inspection of array intensity signals along the genomic axis for candidate CNV regions.

Importantly, higher precision may be possible to achieve for case-parent (trio) studies, as parental array-data can be utilized to inform interpretation of offspring data and *vice versa*. Few methods exist ready for trio-based CNV calling or post-calling assessment from SNP arrays with high specificity of calls and inheritance delineation. Nutsua et al. (2015) exploited offspring-parent CNV call concordance to benchmark CNV callers on a trio cohort, however the approach only used CNV segments coordinates and genomic overlap of those in a manner unaware of the signal intensities, which may be problematic if both offspring and parental calls are artefacts. Scharpf et al. (2012) CNV calling method involves a distance metric between probe intensities in an offspring and a parent. However, it requires a reprocessing of the entire cohort and is not applicable to assessment of already existing calls.

Here, we present SeeCiTe, a post-processing method for trios that assesses evidence for CNVs in an offspring and when possible, suggests inheritance patterns using simultaneously raw signal data from all individuals in the trio. Differently from other methods, SeeCiTe provides intuitive quality categories, a wider range of summary statistics and visual panels displaying raw signal in all three individuals around potential CNVs in the offspring. Thus, SeeCiTe facilitates i) minimization of the number of cases for which visual inspection is needed and ii) visual representations that support inspection with quality classification labels.

Another vexing challenge for the large-scale CNV studies is the need for benchmark datasets enabling quantitative assessment and comparison of CNV calling pipelines. Here we pursue two approaches to develop benchmark data for the evaluation of our tool. The first is based on manual curation of a subset of CNV calls performed by an expert examining signal intensity panels for each potential CNV. The second is exploiting published sequencing-based CNV calls from Genome in a Bottle consortium (Zook et al., 2016).

We use SeeCiTe to refine calls from the Norwegian Mother, Father and Child Cohort GWAS Study (Helgeland et al. (2019) and find that it helps to identify and flag a fraction of CNV calls (including de novo) of lower quality thus increasing the specificity of the calling. The method shows competitive performance with standard filtering strategies while improving on both sensitivity and specificity. The quality categories are well correlated with published sequencing-based CNV sets.

## Methods

### 2.1 Outline

SeeCiTe workflow consists of the following steps (Figure 1), (i) pre-processing of the input; (ii) within-individual summaries calculations; (iii) pedigree analysis;(iv) classification; (v) output generation, where (ii-iv) are the core steps in the method. To maximize the utility of SeeCiTe we recommend to apply it after merging and filtering of initial CNV calls done by PennCNV-trio.

**Fig. 1.**
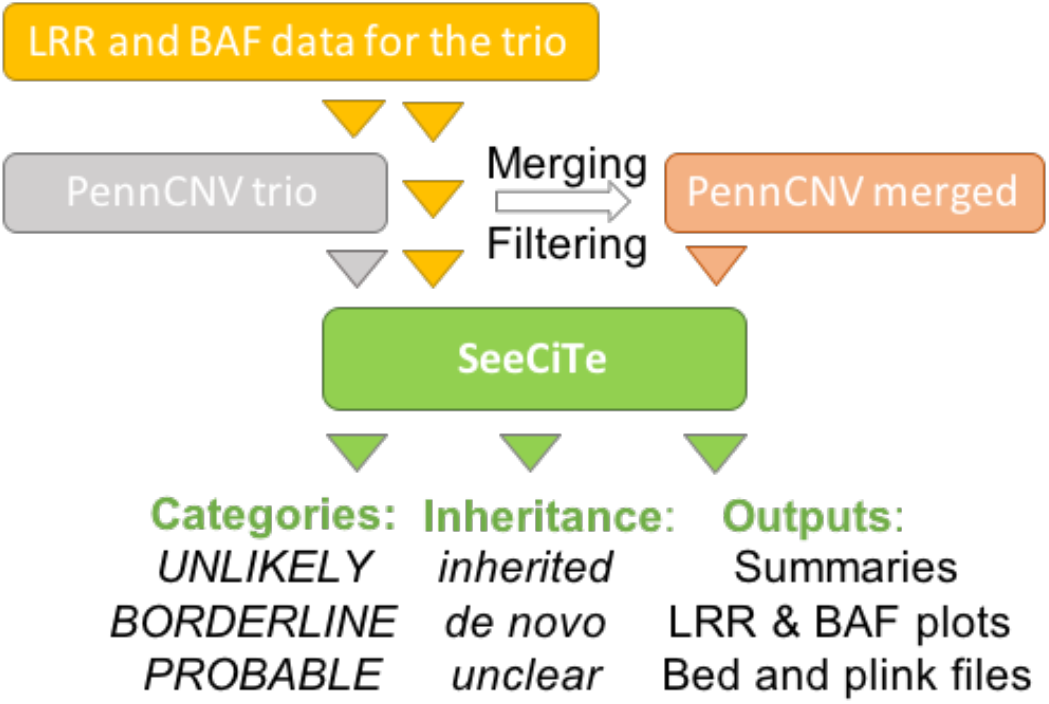
Schematic of the CNV curation for the trio cohort with SeeCiTe. The LRR and BAF signal intensities are used both for initial calling with PennCNV and later for validation. Using PennCNV-trio, typically, a fraction of CNVs will be erroneously split into several fragments that then need to be merged. Filtering by frequency, size and problematic genomic loci is also a standard step to increase specificity of the call set. SeeCiTe then combines all of the inputs to further classify CNV calls, producing output text and pdf files.

#### 2.1.1. Pre-processing: PennCNV inheritance mapping and data points extraction

In the pre-processing phase the original PennCNV-trio (tested with versions 1.0.3 and 1.0.4) calls are mapped onto merged segments and for each CNV, the corresponding set of HMM trio states is parsed into an inheritance status (Wang et al. (2007)), Suppl. Data 1.1. Next, to collect the probe-level information, the raw Log R Ratio (LRR) and B Allele Frequency (BAF) data values are extracted from the GenomeStudio export files for all markers in the region defining the CNV and a flanking area around it (flank), for each individual in a trio using a SeeCiTe wrapper around PennCNV’s infer_snp_allele.pl script. LRR represents normalized total allele intensity at each probe, while BAF is a proxy for A and B alleles ratio, as defined in (Peiffer et al. (2006)). The size of flanks is a parameter set by default to 50 probes, but possible to adjust by the user.

#### 2.1.2. Within-individual summary statistics collection

In the following the standard notation for CNV types is used: del for deletion, dup for duplication, ‘normal state’ for normal copy number state (i.e. CN = 2 for autosomes). LOH stands for Loss of Heterozygosity, which is a necessary but not sufficient condition for calling a del. A region subject to a (one-allele) deletion will only have one allele represented and thus appear as an LOH stretch, but not all LOH regions will correspond to a del.

For each CNV and each individual in the trio, the following summary statistics are calculated (Fig 2A and Fig 3 header):

1. **Allele ratio (BAF)-based summaries** (left top panel on Fig. 2A): CNV type (del, dup or normal state) prediction from the rule-based heuristic classification of the BAF values around the expected allele ratio clusters (Peiffer et al. (2006), detailed in the Suppl. Data 1.2);
2. **Normalized total intensity (LRR)-based summaries** (right top panel on Fig. 2A): Standard deviation of LRR (LRR_SD) in the flanks (up- and downstream pooled together) as a local measure of noise; Number and percentage of LRR values above zero for a del, and below zero for a dup; if this percentage is below a given cutoff (a parameter set to 20% by default) a CNV locus is labelled as supported by LRR; Test whether the LRR values within a CNV locus are shifted from those of the flanks (using the kernel density-based test to reject or accept the null hypothesis that the LRR values in the CNV and in the flanks come from the same distributions, detailed in Suppl. Data 1.3);
3. **Individual-level consistency summary** (middle of the top panel on Fig. 2A): Offspring is consistent with the CNV if both BAF and LRR summaries in 1) and 2) support CNV; A parent is consistent with the normal state if both BAF and LRR support no CNV change, otherwise a parent is consistent with a CNV if BAF supports a CNV;

**Fig. 2.**
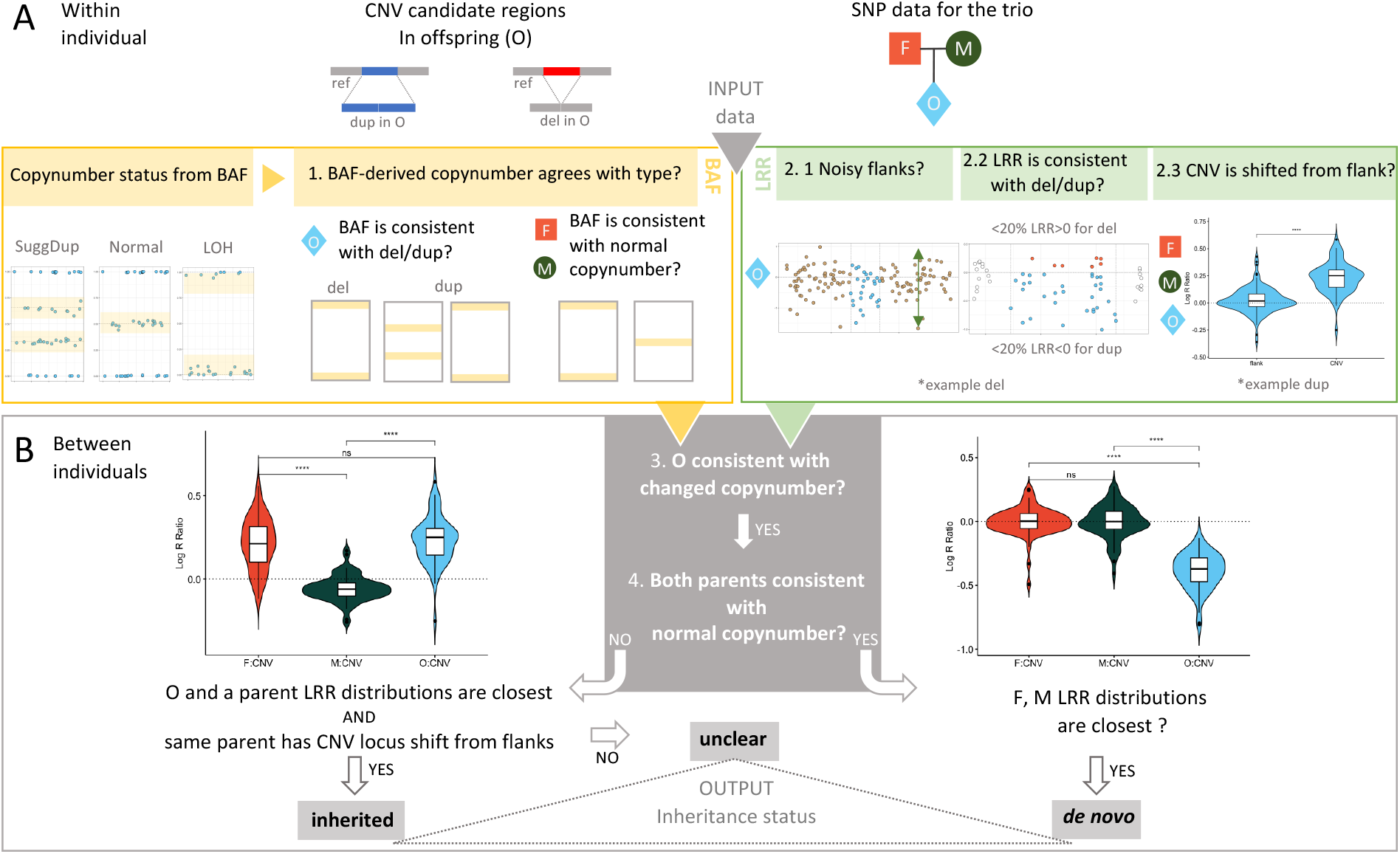
Map of the SeeCiTe method. A) Starting from a candidate CNV call coordinates in an offspring, the method i)extracts LRR and BAF data values for all probes inside the candidate region as well as for flanking 50 probes upstream and downstream (collectively termed as flanks) for each family member – offspring (O), father (F) and mother (M); ii) collects and calculates a set of summary statistics for the locus in each individual; B) The core of the method is direct comparison of the LRR and BAF distributions between the individuals to classify the calls into categories and assign inheritance (left – a dup inherited from the father, right - putative de novo del)

**Fig. 3.**
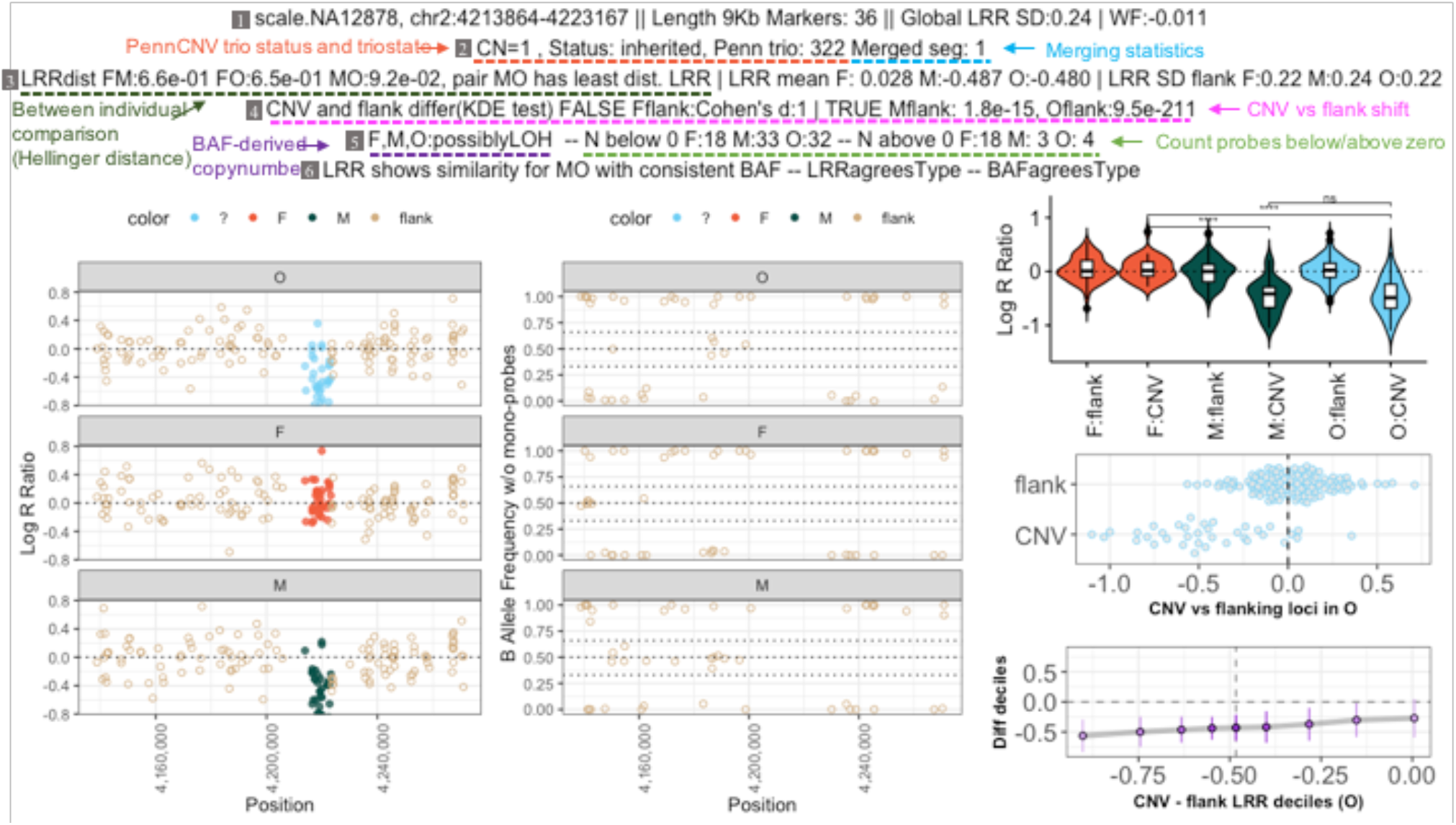
Example of SeeCiTe visual panel for HapMap CEU trio. Header lines numbered for clarity (grey boxes), from top to bottom (1) sample identifier, CNV coordinates, CNV length in kb and probes spanned, genome-wide LRR_SD and waviness factor (WF); (2) PennCNV-trio derived inheritance and original triostate, count of total merged segments (a single segment means the region was not merged as in this example); (3) Between-individuals pairwise LRR comparisons ordered from largest to smallest (Hellinger distance), LRR mean and LRR_SD in the flanks for each individuals;(4)Shift of CNV from flanks as FALSE (no shift detected) or TRUE (with kernel density test p-value);(5)BAF-derived CNV states, counts of probes with LRR values above and below zero; (6) Conclusions of which individuals LRR and BAF distributions appears to be most similar, LRR and BAF consistency for offspring.

#### 2.1.3. Use of pedigree information to assess a CNV call in offspring

The pairwise Hellinger distances (hellinger() in the statip package in R) between LRR distributions for the CNV locus under analysis are calculated for all pairwise combinations of individuals in a trio (Figure 2B). The pair with the smallest distance is recorded as a test result. Then three scenarios are considered:

- *De novo:* When the parents have the smallest Hellinger distance in the CNV locus AND both parents are consistent with normal copy number by BAF, while the offspring is consistent with a CNV (del or dup) (Fig 2B, bottom left).
- Inherited: When a pair offspring-parent has the smallest Hellinger distance for the CNV locus AND both individuals are consistent with a CNV of the same type. (Fig 2B, bottom right);
- Unclear: If none of the above, the CNV inheritance status is labelled unclear.

#### 2.1.4. Quality categorization of the calls in offspring

The final quality classification takes into consideration the input PennCNV trio inheritance and its consensus with the SeeCiTe assignments:

*UNLIKELY*: All putative de novo calls (before or after SeeCiTe inheritance revision) for which LRR_SD value in flank exceeds a given threshold (a value of 0.2 performed well in the datasets in this study)
*BORDERLINE*: When the inheritance assignments from PennCNV and SeeCiTe differ OR contain an ambiguous/unclear label OR when offspring is not consistent with a suggested CNV;
*PROBABLE*: When an offspring is consistent with a CNV AND SeeCiTe inheritance assignment agrees with PennCNV-trio status

#### 2.1.5. Visual panels

For each CNV call in an offspring, SeeCiTe generates a panel of probe-level data for LRR and BAF plotted according to their genomic position for each individual in a trio, for the CNV locus and its flanking regions. Summary statistics are condensed in the header for all three individuals. Boxplots (for all individuals) and decile plots (for the offspring) for LRR are visualized. (Figure 3)

### 2.2 Evaluation on the real cohort and public HapMap trio data

#### 2.2.1. MoBa cohort

The Norwegian Mother, Father and Child Cohort Study (MoBa) is a population-based pregnancy cohort study conducted by the Norwegian Institute of Public Health. (Magnus et al. (2016)) Participants were recruited from all over Norway from 1999 to 2008. The women consented to participation in 41% of the pregnancies. The cohort now includes 114,500 children, 95,200 mothers and 75,200 fathers. The current study is based on version 9 of the quality-assured data files released for research. The establishment of MoBa and initial data collection was based on a license from the Norwegian Data protection agency and approval from The Regional Committees for Medical and Health Research Ethics (#2012/67). Blood samples were obtained from both parents during pregnancy and from mothers and children (umbilical cord) at birth.

#### 2.2.2. MoBa data pre-processing

Two batches of MoBa trio samples were processed in the current study, MoBa1 was assayed on Illumina’s HumanCoreExome-12 v.1.1 (MoBa1.12) and HumanCoreExome-24 v. 1.0 (MoBa1.24) and MoBa2 was genotyped using the Illumina’s Global Screening Array v. 1.0 as described in Helgeland et al. (2019). CNVs were only called in samples that had passed SNP-based genotyping and quality control. Full details on CNV calling and filtering prior to SeeCiTe can be found in Suppl. Data 2.

#### 2.2.3. Public HapMap CEU trio

Publicly available CEL files for HapMap CEU trio (NA12878, NA12891, NA12892) were obtained from ftp://ftp.ncbi.nlm.nih.gov/hap-map/raw_data/hapmap3_affy6.0/. We used PennCNV-Affy module to generate the signal intensity data for two replicas of the trio assay (Tesla and Scale), suitable for CNV calling. The data was processed as in Suppl. Data 2, except for frequency filtering, which was not done on the HapMap trio data.

#### 2.2.4. Benchmark datasets

Two manually curated datasets were produced for MoBa and one for public short-reads sequencing based data:

1. Of the SeeCiTe output classification categories of MoBa1.24, all UNLIKELY and BORDERLINE calls, as well as all de novo calls were examined visually by an experienced clinical laboratory geneticist using LRR intensity and BAF plots of the trio, resulting in inspection labels for each CNV, either “bona fide” for likely real CNVs or “noise” for highly likely artefacts.
2. The above was repeated for MoBa2. Furthermore, for the approximately 2000 probable inherited calls in the MoBa2 set, visual inspection was performed on 180 calls, randomly obtained using the same distribution of events per chromosome as in a de novo call set for visual inspection. Additionally, some calls on the extremes of quality parameters were examined: a) 10 inherited CNV events in which the offspring had high fraction of probes in direction discordant with the copy number predicted; b) 10 CNV events with the highest LRR_SD in flanks.
3. We used high confidence published golden standard CNV calls from 1000 Genomes (GS1) (Sudmant et al. (2015)) and (Parikh et al. (2016)) (GS2). The two sets were combined as follows: 1) at least 80% reciprocal overlap of GS1 and GS2 resulting in a more stringent set, Ref_intersection (1540 dels, 0 dups); 2) the union of GS1 and GS2, Ref_union (3021 dels, 78 dups/ins). As only deletions are available in the intersection set, we used only deletion calls for the sequencing-based benchmark.

#### 2.2.5. CNV quality control strategies tested

Using the benchmark data sets defined above, we compared SeeCiTe with other frequently used methods (Table 1). The details of how the CNVs and their scoring between the methods were matched (when possible) are expanded in Suppl. Data 3.1.

**Table 1.** Methods and quality control strategies used for CNV filtering

| Method | Short name | Quality categories | Reference |
| --- | --- | --- | --- |
| PennCNV merged | PennCNV | Confidence score (& size) | Wang et al. (2007) |
| Quality Score for PennCNV merged | QS | Probability of being detected by other method | Mace et al. (2016) |
| PennCNV-trio merged (iterative) | PennCNV-trio | Frequency & size | Wang et al. (2007) |
| SeeCiTe | SeeCiTe | *UNLIKELY, BORDERLINE, PROBABLE* | Current study |
| Consensus | Consensus | Intersection of PennCNV and QuantiSNP | Common practice |

#### 2.2.6. ROC curves

The Receiver Operating characteristic (ROC) and Precision-Recall curves and Area Under the Curve (AUC) values were calculated using an R package precrec (Saito & Rehmsmeier, 2017) on the benchmark dataset MoBa1.24.

#### 2.2.7. HapMap comparisons

Due to the restricted resolution of the array, we limited the comparisons to deletion calls that span > 20 kb (> 10 probes) in both reference segments and arrays. To compare array calls with the reference we considered genomic overlap of the two sets. A CNV was tagged as None if zero overlap existed between an array call and any of the (size filtered) reference sets; Concordant in case of the reciprocal overlap of at least 50% and Low_overlap otherwise.

## Results

### SeeCiTe software

SeeCiTe is implemented as an R package (developed in R version 3.5.1) and is freely available at https://github.com/aksenia/SeeCiTe, distributed under the MIT license. Current implementation is tailored to native PennCNV output formats, while the final classified data is written in generic UCSC bed and plink data types. The software comes with a tutorial, documentation and example data from the public HapMap repository.

### CNV refinement with SeeCiTe in the Norwegian Mother, Father and Child Cohort Study

We analyzed two trio data sets from MoBa (MoBa1 and MoBa2, Table 2). MoBa1.12 was used for parameter calibration while MoBa1.24, MoBa2 and a publicly available CEU HapMap trio were used to validate and assess the performance of the tool.

**Table 2.**
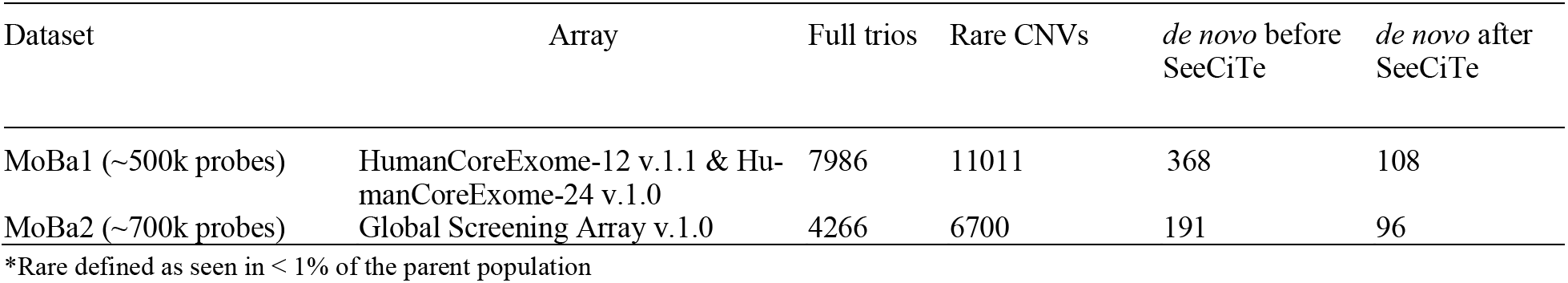
Statistics of CNV calling in MoBa cohorts. Left to right, Dataset and resolution; Array used; Count of full trios in the core set of Norwegian ancestry; Count of CNVs after the frequency and size filtering according to Suppl. Data 2.3; Count of de novo CNVs before and after SeeCiTe.

### Evaluation of SeeCiTe

#### 3.3.1 Assessing SeeCiTe performance on the benchmark datasets

We assessed SeeCiTe quality classification by calculating the percentages of bona fide calls in each examined category for the benchmark set from MoBa2 (see 2.2.4).

SeeCiTe classified candidate CNVs in good concordance with the benchmark (expert assigned) labels (Table 3), ranging from the *PROBABLE* category, as expected, capturing the majority of bona fide calls (94-98%), to the *BORDERLINE* category with less bona fide calls and the *UNLIKELY* category that was enriched in artefacts (only 4-5% bona fide calls).

**Table 3.** Ratio of bona fide (true) calls for each SeeCiTe quality category in MoBa2 benchmark. Counts and percentages of manually classified bona fide calls in each inspected SeeCiTe category – all inspected calls, only *de novo* calls.

| CNV category | *UNLIKELY*<br>Fraction (True/total) | *BORDERLINE*<br>Fraction (True/total) | *PROBABLE*<br>Fraction (True/total) |
| --- | --- | --- | --- |
| All inspected | 4 % (3/70) | 83% (138/167) | 98% (285/290) |
| *de novo* only | 5% (3/54) | 33% (3/9) | 94% (85/90) |

All 200 inspected putatively inherited calls classified as bona fide inherited CNVs, which is expected due to their double support from two individuals (Suppl. Fig S3A and Suppl Data 3.2). SeeCiTe likelihood categories closely followed the trends of LRR intensity variation (LRR_SD) in flanks. (Suppl. Fig S3B for separation by local versus global LRR_SD).

#### 3.3.2 Comparing the accuracy of SeeCiTe with other methods

We next compared SeeCiTe with other PennCNV pipelines and a Consensus method (Table 1) on the benchmark dataset MoBa1.24 (see 2.2.4). The relationships between PennCNV-based pipelines considered are shown on the Suppl. Fig S2. SeeCiTe is a direct downstream step of PennCNV-trio in which the quality control step included filtering by frequency and size (Suppl. Fig S2). The majority of the calls in the benchmark MoBa1.24 (84.3%) were classified as *PROBABLE* inherited calls, and as section above demonstrated, were high confidence calls, on which PennCNV-trio and SeeCiTe methods agree. However, 2.5% of the total calls in PennCNV-trio in this set were labelled as “noise” in our benchmark set and could be potential artefacts. In SeeCiTe categories, the majority of calls identified as noise were in the *UNLIKELY* category (84%), while 7.6% were in *BORDERLINE* and only 1.4% (two calls) in the *PROBABLE* category. This shows that applying the SeeCiTe tool to post-process the output from PennCNV-trio aided in identifying likely false positive calls, while losing few high-quality CNVs.

For the purpose of this analysis, the Consensus method was implemented using the calls in the intersection of PennCNV and QuantiSNP. We defined a call as consensus if any probes overlapped between the two callers. With this definition, the agreement between SeeCiTe and Consensus methods was high (Suppl. Data 3.3). However, the Consensus method was overly conservative as it rejected 53 inherited calls (Suppl. Fig. S4A) which upon additional manual inspection were validated as bona fide inherited calls. This behavior of the consensus methods is well known (Pinto et al., 2011).

Next, we compared the quality score distribution for the methods that provide a score. As SeeCiTe does not report a single unified quality score (rather relying on several discrete and continuous variables in the decision process) we used the LRR_SD in flanks as the best contiguous predictor of CNV call quality in SeeCiTe. For all four methods the overall score distributions had an apparent peak for noise, but the ability to separate between the bona fide calls and noise varied, with SeeCiTe proxy score and QS performing better than two other methods. (Suppl Fig. S5A). Note that, if following recommendation of Mace et al. (2016) on the cutoff of 0.5 for QS, one would lose in sensitivity without gaining in specificity, as the discriminative value of the score was at its highest at around 0.15 in this dataset. We further assessed all methods using ROC and Precision-Recall to calculate the area under the curve on the balanced subset of 166 bona fide and 111 noise labelled CNV calls of the benchmark set MoBa1.24 (Suppl. Fig S5B-C). This also confirmed SeeCiTe representative score as the top performer in separating the two categories with most optimal specificity to sensitivity trade off.

#### 3.3.3 Validation in HapMap CEU trio

To complement our assessments of the SeeCiTe method on an independent dataset with an existing set of sequencing-based CNV calls, we used publicly available data for the HapMap CEU trio NA12878, NA12891 and NA12892.

We ran SeeCiTe on publicly available Affymetrix6.0 array data for two replicas of this trio (“Tesla” and “Scale”) and manually curated the calls using visual panels, independently for each sample. Of note, the two replicas varied in signal quality with Tesla having larger global LRR_SD of 0.28 as compared to 0.24 of Scale. This provided an opportunity to investigate to what extent errors, due to noise, can be made in the curation process using visualization panels. We did this by assessing the concordance between the curation results obtained for the two samples and the concordance with the sequencing-based benchmark set.

##### Between-replicates comparison

At a size cutoff of 20kb, we detected 29 and 24 deletion calls in Scale and Tesla samples respectively. Among these, 18 calls were shared between the samples (14 pairs with exact breakpoints match and 4 pairs with varying breakpoints). The 18 pairs of calls shared between the samples had higher concordance with the benchmark set segments compared to singleton calls, e.g. only 41% (7/17) singleton calls overlapped a benchmark segment (in the Ref_union only), as opposed to 72% (26/36) of shared calls (12 of which overlapped segments in the Ref_intersection set). Upon visual inspection, 8.3% (3/36) of shared and 35.3% (6/17) of singleton calls were flagged as noise, with more noise in Tesla compared to Scale (Fig. 4).

**Fig. 4.**
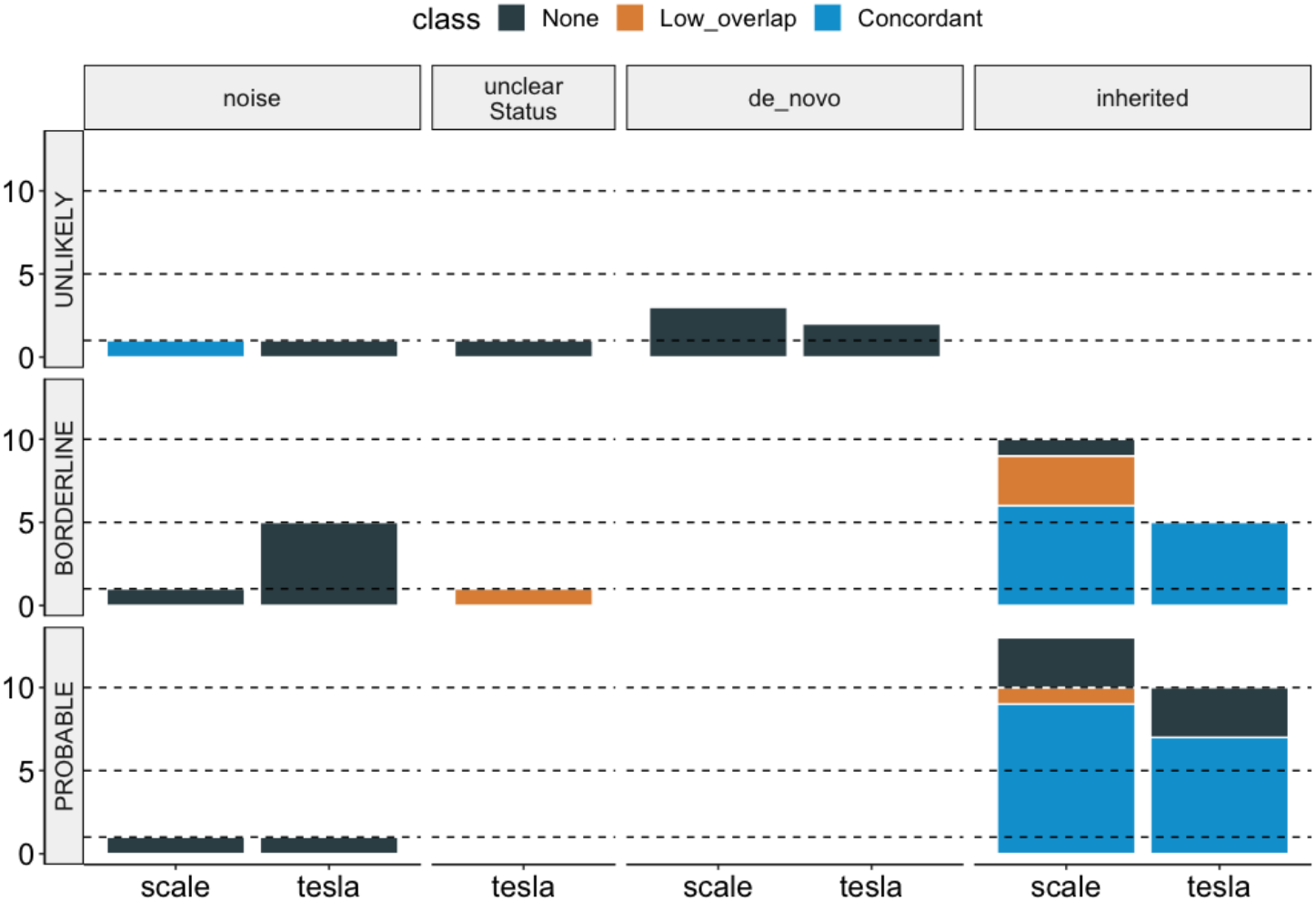
Comparison of SeeCiTe-assisted HapMap sample classification to a published benchmark. Counts of CNV calls in each replicate binned by concordance with the reference (color coding: None for no overlap with any of the reference; Low overlap – less than 50% reciprocal overlap with a benchmark set deletion; Concordant >=50% reciprocal overlap with a reference deletion), expert curation labels (top panels); SeeCiTe classification (left panels), unclearStatus means CNV in an offspring but inheritance is not clear due to noisy signal in parents

The majority (16/18) shared call pairs were concordant in independent manual inspections, except for two, for which the difference stemmed from local lower quality of the signal in the father that prevented PennCNV trio and following visual assessment to resolve the inheritance unambiguously.

##### Comparison to the benchmark set

SeeCiTe quality classes were roughly in agreement with benchmark concordance, with *UNLIKELY* calls having the least of overlaps and *PROBABLE* calls with the highest fraction of Concordant calls. Notably, Scale, a less noisy sample, had a higher fraction of Concordant calls (16 vs 14) and lower fraction of calls with no overlap (9 vs 11) (Fig. 4).

To conclude, we observed a high degree of concordance of SeeCiTe likelihood assignments with the benchmark deletions set. The errors that were made with both PennCNV trio method and visual assignment based on the signal panels, stemmed from the intrinsic noise-to-signal ratio and could be mitigated, to a large extent, by controlling flanking LRR_SD as well as filtering by known common CNV polymorphic loci.

## Discussion

In this study we presented and evaluated a method that exploits Mendelian inheritance patterns to curate and aid interpretations of CNV calls in offspring-parents trios. For each of the CNV calls, SeeCiTe performs analyses to assess how well the call is supported by the underlying data resulting in labelling each call as either *UNLIKELY, BORDERLINE* or *PROBABLE*. This provides an intuitive classifier that guides attention to a much smaller subset of calls that ought to be examined more in detail. To aid the examination we devised a visual panel of local LRR and BAF intensities for a trio together with key quality summaries. We show that together with the CNV-quality labelling these panels serve as a good proxy for the ground truth. We further demonstrate on a public dataset of HapMap CEU trio that SeeCiTe quality categories are in good concordance with the published sequencing-based CNV sets.

We are not aware of another method that provides such in-depth examination, summaries and convenient visual panels as SeeCiTe. SeeCiTe method is developed to be used downstream of PennCNV, which is a stable, commonly used and contiguously maintained software suite. We did not evaluate the workflow on other callers and cannot in principle guarantee the same performance.

The over-merging is a potential issue with the CNV segments merging procedure, solely based on distances between the calls. Depending on the priorities of each analysis, the parameters of merging can be adjusted, but generally, the over-merging issue is mitigated by SeeCiTe, since LRR distribution assessment in the method will inevitably highlight calls that have too many probes with normal state intensities. Notably, using visual inspection, we found complex CNV events that are inherited as a haplotype block of two dups/dels and a normal state between, which we find reasonable to count as a single event, but we are not aware of any caller that is able to make this judgement.

We find that putative de novo calls have higher false positive rates compared to putative inherited calls. This may not be surprising, as inherited calls are supported by two events, unlike de novo calls. In SeeCiTe this asymmetry is addressed by implementing a hard cutoff on the value of LRR_SD in the flanking regions of putative de novo calls, thus tagging calls with too high noise as UNLIKELY, recommending them to be inspected. In general, due to the low proportion of de novo calls, it is feasible to inspect all potential de novo candidates and this is what we recommend.

The LRR_SD in flanks is useful as a proxy score for SeeCiTE, as illustrated by the ROC analysis. However additional measures, that also verify the intensity level and direction of the shift of LRR intensities in the CNV locus, increase the specificity (data not shown).

We used visual panels for benchmarking due to the lack of the ground truth in the cohorts in this study. Even though we complemented evaluation with a public golden standard set in HapMap controls, we note that the cell-line derived HapMap data do not compare fairly to MoBa studies based on whole blood samples due to intrinsic biases and differences of the two types of samples (Joesch-Cohen & Glusman, 2017).

For further work, the parameters identified in the current study can be in principle used in an unsupervised machine learning workflow to provide even better automated discrimination of CNV calls likelihood. Such workflow would incorporate simultaneous calibration of all parameters involved which would facilitate more in-depth understanding of the contribution of each variable and their interdependence.

To summarize, in this article we presented the SeeCiTe method that helps to refine and assess the CNV calls in offspring for large-scale trio studies. The tool simultaneously provides quality categorization and visual panels for each CNV call. Our analyses demonstrated that the method has good performance on several types of arrays and good concordance with benchmark data including calls based on sequencing. As such, SeeCiTe allows to streamline quality control and manual steps often performed when utilizing array data to call CNVs in large scale trio cohorts.

## Data Availability Statement

The data underlying this article were provided by the Norwegian Institute of Public Health under license from the Norwegian Data protection agency and approval from The Regional Committees for Medical and Health Research Ethics (#2012/67). Access to genotype data can be obtained by direct request to the Norwegian Institute of Public Health (https://www.fhi.no/en/studies/moba/for-forskere-artikler/gwas-data-from-moba/).

## Acknowledgements

The Norwegian Mother, Father, and Child Cohort Study is supported by the Norwegian Ministry of Health and Care Services and the Ministry of Education and Research. We are grateful to all the participating families in Norway who take part in this ongoing cohort study. We thank the Norwegian Institute of Public Health (NIPH) for generating high-quality genomic data.

## Contributions

KL called CNVs and performed initial QC and filtering; KL designed, implemented and tested the method. ØH obtained the data permissions, performed SNP-based genotyping, QC and ancestry analyses prior to CNV work. PRN and SJ provided funding and access to the data, SJ conceived the study and together with IJ supervised all aspects of the work. KL, SJ and IJ wrote the manuscript with contribution from all authors.

