## Supplementary data for "SeeCiTe: a method to assess CNV calls from SNP arrays using trio data"

**Supplementary information**

### **1. Details of the SeeCiTe algorithm**

#### **1.1. PennCNV-trio inheritance mapping**

**The trio state for each CNV call was first disambiguated by collapsing the redundancies (e.g. “232-232” to “232”) followed by definition of inheritance status based on total copynumber (Table 1 in (Wang et al., 2007); 0,1 for a deletion, >2 for duplication) predicted in each individual in a trio, as inherited/*de novo* or ambiguous in case of conflicting patterns. If all original CNV calls within a merged segment had the same status, it remained the final assignment, otherwise if one of the calls had an ambiguous status, or if the inheritance was discordant within a merged segment, an ambiguous status was assigned.**

#### **1.2. Inferring CNV type from B Allele Frequency clusters**

Normalized BAF clusters were used to infer the underlying copynumber relying on the expected values for each copynumber (**Suppl. Fig S1**). The margins around the clusters and the choice of informative cluster centers were calibrated empirically on the subset of MoBa1.12 and proved efficient for the rest of the data. With the definition of clusters as in **Suppl. Fig S1**, the following variables were defined:

- **het -** count of probes representing the normal (autosomal) copynumber CN = 2;
- **center** – count in a more stringent variant of the heterozygous cluster;
- **middle**– count everything except the loss of heterozygosity points;
- **dup** – count of probes in the largest among cn3, cn4 or cn5 (CN=3, 4 or 5; Suppl. Fig. S1);
- **dup total** – count of probes in the **middle** minus those falling in **het**.

After that the rules outlined in Suppl. Table 1 below were applied to infer a CNV type:

**Supplementary Table 1. Rules for inferring CNV type based on BAF**

| **Loss of Heterozygosity** | **Duplication** | **Normal state** |
| --- | --- | --- |
| fraction (middle) < 0.05 *AND*  count (het + cn3) < 3  *OR*  count (het + dup) < 2 | count (het) < count (dup total) *OR*  *IF* count (het) == count (dup total) *AND*  ((dup = cn4 *AND* count (dup) >1) *OR*  (dup = cn5 *AND* count (dup) > 1) *AND*  fraction (dup) > 0.25 *AND*  count (center) < count (dup)) | count(het) > count (dup total)  *OR*  fraction (dup) < 0.1 |

### **1.3. Testing whether the LRR distribution in a CNV locus is shifted from those of the flanks**

To mitigate the false positive significant differences between the LRR in CNV and flanking regions distributions we implemented a two-step procedure. **First, a Hedges effect size g was calculated between the two distributions (R function cohen.d(…,hedges.correction = T), and any value below the cutoff value 3 was considered as no difference between the two distributions . If the g value exceeded the cutoff of 3, a kernel density-based test was performed on the two distributions employing a cutoff p-value of 0.01 (kde.test in from ks package in R). Thus, a pair CNV and flank could be declared not different either following a first step (small Cohen’s distance) or an actual distributions shift test with a p-value cutoff of 0.01.**

### **2. Cohort data processing prior to SeeCiTe**

#### **2.1. CNV calling**

The Log R Ratio (LRR) and B Allele Frequency (BAF) values were extracted using GenomeStudio (version v.2011.1 for MoBa1 and v.2.0.3 for MoBa2) (<https://www.illumina.com/techniques/microarrays/array-data-analysis-experimental-design/genomestudio.html>). **CNVs were called with PennCNV (Wang et al. (2007),** version 1.0.3 for MoBa1 and 1.0.4 for MoBa2**) in genome build** GRCh37/**hg19 with GC-waves adjustment, using default parameters for first-pass quality control (LRR_SD < 0.3, BAF_drift < 0.01, |WF| < 0.05) and merging. Samples with more than 100 (130) calls for MoBa1 (Moba2) were removed from further analyses. PennCNV-trio was run after PennCNV to refine the calls in trios. PennCNV trio module uses Bayes posterior probabilities to assess a copy number state in each member of a trio in a given locus. The software may add calls that were missed during initial calling as well as redefine the boundaries within a trio. QuantiSNP (Colella et al. (2007)) was used to call CNVs in MoBa1.24 to assess consensus calling. Default recommended parameter values (Log Bayes Factor < 30) were used to define quality passing CNV calls.**

#### **2.2. Iterative merging of fractionated CNV calls**

Given that the size filtering is a necessary step in maximizing specificity of the CNV call set, it is crucial to minimize the number of CNV events that are fractionated. The PennCNV suit contains a utility script **clean_cnv.pl** that merges two consecutive calls having the same copy number state depending on the size of the gap between the two calls. The threshold is calculated as a percentage (default 20%) of the region encompassing the given calls (calculated in base pairs or probes) (Fang and Wang (2018)).**We applied an iterative merging procedure that involves repeated application of the merging utility, alternating between gap fraction calculations from percentage of probes and percentage of basepairs (-bp flag) starting from gap at 50%, down 10% in each cycle, with output of each cycle fed as input to the next, until no further merging was happening. In the current dataset, the maximal merging observed included six cycles.**

#### **2.3. CNV set filtering**

**Only the trios in which all individuals passed the default PennCNV quality control (LRR_SD < 0.3, BAF_drift < 0.01, |WF| < 0.05) were selected for further analysis. Common CNVs were removed using a 1 % allele frequency threshold as defined among parents with PLINK v1.07 (Purcell et al. (2007)). Known problematic regions, such as centromeres, telomeres (UCSC Table Browser,** NCBI37/hg19 human genome assembly, “gap” table**) and immunoglobulin loci were removed. Only CNV calls spanning >10 probes and >100kb we retained, based on the resolution of the chips in the MoBa datasets.**

### **3. Evaluation of SeeCiTe**

#### **3.1. Matching the various pipelines**

The datasets processed with (PennCNV), (QS) and (Consensus) did not undergo the extensive merging and thus could contain multiple calls that correspond to a single call in (SeeCiTe). To account for that we matched each call in SeeCiTe with all calls from (PennCNV /QS/ Consensus) that were either identical or contained in a given call from SeeCiTe. In case of multiple calls, we calculated a median of the quality scores for matching calls for each method. The quality score in PennCNV represents the log likelihood of the most likely state minus the second most likely state in the PennCNV Markov chain model for a given CNV call. For the convenience of comparison here, we log transform the original confidence score. The quality score QS in Mace et al. (2016) represents a probability of a call to be a consensus call (e.g. to be called by another software), resting on a reasonable assumption that calls identified by several software packages are more likely to be real calls. This is achieved by using a linear regression model trained on an independent dataset by Mace et al. (2016) and supplied with the software. The Consensus method in its basic form does not have a score but rather a binary inclusion-exclusion variable based on whether a call is found in the intersection of the two callers or not (Consensus_binary). Since QuantiSNP provides a score - a Log Bayes Factor of most probable copy number state (maximum of Factors for all possible states), we also implemented a variation Consensus_score with log transformed original score and a zero whenever a call was not a consensus.

#### **3.2. Details of misclassified calls by SeeCiTe in section 3.3.1**

Among the three putative *de novo* calls assigned to the *UNLIKELY* category that were recognized as *bona fide* by visual inspection, two had intensity variation in the flanks close to the cutoff values used and one had too many probes with intensity value inconsistent with the predicted copy number, and could potentially be a mosaic event in a child. This explained and justified a conservative placement of these calls in the *UNLIKELY* category by SeeCiTe.

Out of 167 borderline quality calls, 5 calls (3%) were marked as over-merged e.g., upon visual examination we observed stretches of normal copy-number state between the CNVs, which clearly contributed to the shift of LRR distributions towards borderline values. Notably, we observed a clear bias in the borderline inherited calls, as opposed to randomly drawn “representative” probable calls, in that the parental values had less noise in flanking regions than those of the offspring. (Suppl. Fig. S3A) This is an example of an additional support that Mendelian inheritance patterns lend to likelihood ascertainment.

Of 90 *PROBABLE* *de novo* calls (e.g. passing SeeCiTe quality tests and both PennCNV-trio and SeeCiTe agreeing on an inheritance status), only 3 calls (3.3%) were identified as artefacts and one call (1.1%) as likely, but inherited. Upon inspection of this misclassified inherited call, it is clear that the boundaries of the initial raw CNV call by PennCNV-trio were wrong and included three parts: a segment, likely inherited from the father, a segment with normal state and potential *de novo* duplicated segment.

#### **3.3. Details of SeCiTe and Consensus method errors**

The misclassification of *bona fide* calls is similar between the two methods, with SeeCiTe having slightly lower error rate (3% false negatives against 5% in consensus). The two calls that were labeled as *PROBABLE* in SeeCiTe, but on a close inspection turned out to be less trustworthy, are relatively small calls (<20 probes) with higher than average GC-content in the locus (0.013 and 0.017 versus median of 0.0076). Interestingly, there are 3 shared calls between the false negatives (in total 5 in SeeCiTe and 9 in consensus), indicating overall problem with these loci rather than particular method. Of the remaining two false negative calls for SeeCiTe, they are large (>300 probes) and thus can tolerate more noise in the flanking regions. The remaining 6 calls that were wrongly rejected by the consensus approach were classified as *bona fide* *de novo* upon inspection.

**SeeCiTe: a method to assess CNV calls**

**from SNP arrays using trio data**

**Supplementary Figures**

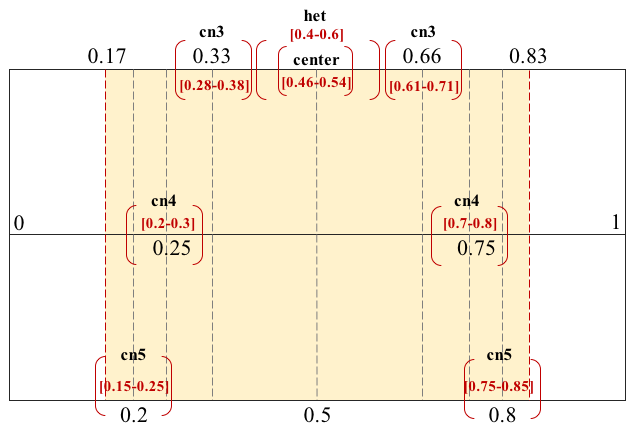

###### **Suppl. Fig S1. Definition of B Allele Frequency (BAF) clusters used for inferring a CNV type.** BAF represents normalize ratio of alleles A and B. The expected values of BAF for i) normal state (eg. heterozygous, no CNV) is around 0.5 – het cluster at 0.5+/-0.1 and het center cluster at 0.5+/-0.04; ii) loss of heterozygosity is manifesting in values at only around 0 and 1 – the intervals [0,0.17] and [0.83,1]; iii) duplications are characterized by different clusters, depending on a copy number, e.g. copynumber 3, upper line (cn3) at 0.33+/-0.05 and 0.66+/-0.05, copynumber 4 (cn4), middle line, at 0.25+/-0.05 and 0.75+/-0.05, copynumber 5 (cn5), lower line, at 0.2+/-0.05 and 0.8+/-0.05

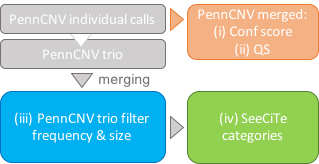

###### **Suppl. Fig S2. Relationship between PennCNV-based pipelines.** SeeCiTe method is applied downstream of PennCNV trio, iterative merging and frequency and size filtering thus sharing exact CNV boundaries with (c), while PennCNV merged (b and d) is an independent default merging that may be a more fractionated call set

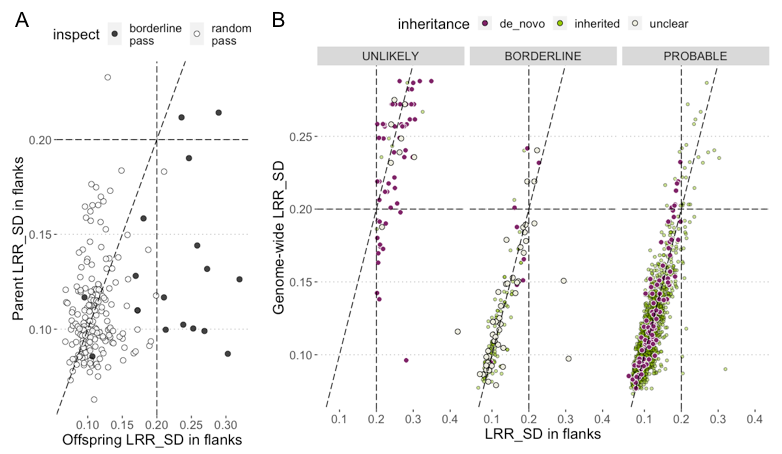

###### **Suppl. Fig. S3. Characteristics of LRR_SD distribution in benchmark set MoBa2.** A) 200 inspected inherited calls (180 drawn at random and 20 borderline quality), with offspring LRR_SD values in flanks on x-axis and that of the parent of origin on the y-axis; B) Distribution of LRR_SD values genome-wide per sample (y-axis) and in regions flanking each CNV call (x-axis) for the three quality categories: *UNLIKELY, BORDERLINE, PROBABLE,* colored by inheritance assigned by the method

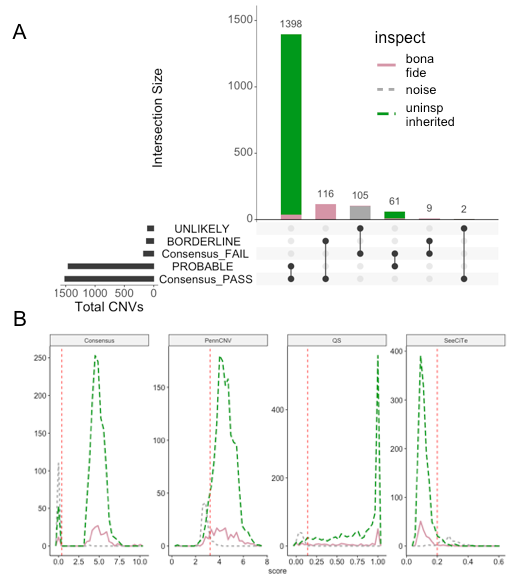

###### **Suppl. Fig S4. Comparison of SeeCiTe categories and other methods.** A) Pairwise intersection of SeeCiTe classes (*PROBABLE, BORDERLINE,UNLIKELY*) and consensus method – Consensus_PASS/Consensus_FAIL depending on whether a call was found by both callers of just one; B) Comparison of Consensus, PennCNV, QS and SeeCiTe in terms of separation of bona fide and noise labeled calls represented as the density of the calls with score range on the x-axis;. Vertical dashed line indicates hypothetical cutoff on the quality score for each method; The uninspected inherited calls are likely good CNV calls and their score distribution follows closely that of *bona fide* calls

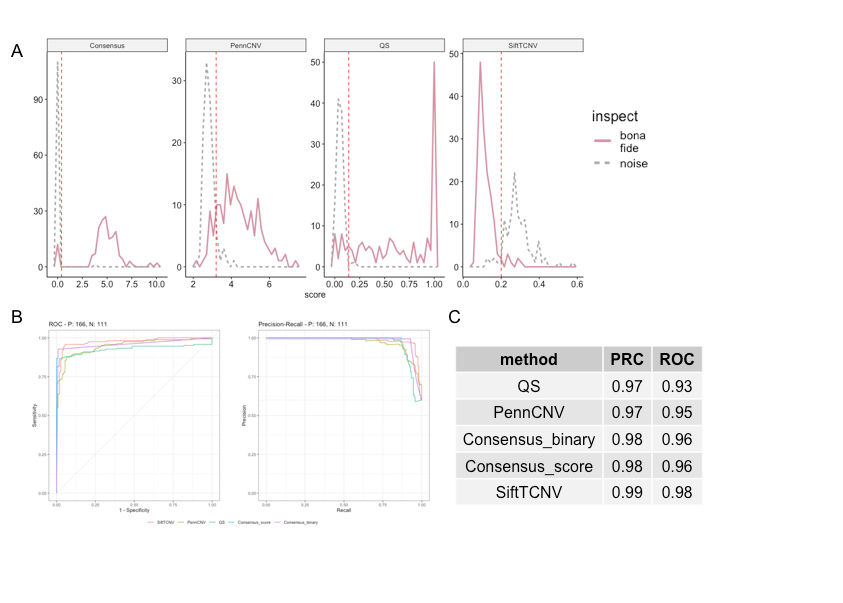

###### **Suppl. Fig S5. Comparison of SeeCiTe categories and other methods**. A) Comparison of Consensus_score, PennCNV, QS and SeeCiTe in terms of separation of bona fide and noise labelled calls represented as the density of the calls with score range on the x-axis; Vertical dashed line indicates hypothetical cutoff on the quality score for each method; B) ROC (left) and Precision-Recall (right) curves calculated on the MoBa1.24 dataset bona fide and noise classifications; C) Table of AUC values for both ROC and Precision-Recall, sorted increasingly, with SeeCiTe showing the best values for both
